# Selective ablation of Nfix in Macrophages preserves Muscular Dystrophy by inhibiting FAPs-dependent fibrosis

**DOI:** 10.1101/2021.05.12.443809

**Authors:** Marielle Saclier, Giulia Temponi, Chiara Bonfanti, Graziella Messina

## Abstract

Muscular dystrophies are genetic diseases characterized by chronic inflammation and fibrosis. Macrophages are immune cells that sustain muscle regeneration upon acute injury but seem deleterious in the context of chronic muscle injury such as muscular dystrophies. Here we observed that the number of macrophages expressing the transcription factor Nfix increases in two distinct murine models of muscular dystrophies. Plus, we showed that the deletion of Nfix in macrophages in dystrophic mice delays fibrosis establishment and muscle wasting until 6 months of life. Indeed, macrophages lacking Nfix express more TNFα and less TGFβ1 thus promoting apoptosis of fibro-adipogenic progenitors. Moreover, pharmacological treatment of dystrophic mice with ROCK inhibitor accelerates fibrosis through the increase of Nfix expression by macrophages. Thus, we identify Nfix as a macrophage profibrotic actor in muscular dystrophies, whose inhibition could be a therapeutic way to rescue the dystrophic disease.

## Introduction

Muscular dystrophies (MDs) are a heterogeneous group of inherited disorders caused by mutations in genes coding for the structural dystrophin-glycoprotein complex inducing its disassembly. This leads to sarcolemma fragility and myofiber necrosis especially during intense contractile activity, provoking cycles of regeneration-degeneration^1–3^. These continuous cycles induce muscle wasting, chronic inflammation, fibrosis, which in the most severe cases can lead to patients’ death^4, 5^. MDs remain incurables and until now, only drug therapy brings some mild results. Indeed, glucocorticoids uptake delays MDs progression but has numerous side effects such as bone fragility, weight gain, mood change but also muscle weakness when taken chronically^6–9^.

Dystrophic muscle cells are at the origin of the disease, but several other cell types participate in MD progression and muscle loss. Macrophages (MPs) are associated with fibrosis establishment particularly toward their action on fibro-adipogenic progenitors (FAPs)^10, 11^. While timely specific populations of MPs are required for successful muscle regeneration upon acute injury, it is not clear if MPs act negatively or positively in MDs. It seems that a general impediment of MP infiltration within *mdx* muscle, the murine model for Duchenne Muscular Dystrophy (DMD), improves both muscle histology and function^12–14^. But in some studies, the decrease of MP infiltration is not correlated with improvement of MDs. Indeed, *mdx* mice deleted for urokinase plasminogen activator display an increase of degenerative area and fibrosis together with a decrease of muscle function that correlates with a decrease of macrophage infiltration^15^. More recently, a study demonstrated that a decrease of MP infiltration induced by blood monocyte depletion promotes the conversion of satellite cells (SCs), the muscle stem cells, into adipocytes^16^. The anti-inflammatory TGFβ protein, which is mainly secreted by MPs, induces extracellular matrix (ECM) component expression by FAPs and thus promotes fibrosis^17–19^, but conversely, pro-inflammatory markers have been associated with pro-fibrotic function of MPs and DMD worsening^20–22^. More than the presence or absence of MPs, it seems that MPs’ phenotype and specific cytokines secretion play an important function in MDs progression.

Nuclear factor I X (Nfix) is a transcription factor belonging to the highly conserved DNA-binding nuclear factor one family (Nfi) together with Nfia, Nfib and Nfic^23^. We previously demonstrated that Nfix is essential to muscle development by driving the transcriptional switch from embryonic to fetal myogenesis^24^ and, in the adult, SCs lacking Nfix failed to timely differentiate inducing delay of muscle regeneration upon acute injury^25^. Importantly, the deletion of Nfix in two murine models of MDs improves muscle histology and function by delaying muscle regeneration, demonstrating the concept that preserving muscle from cycles of degeneration-regeneration is more beneficial than boosting muscle regeneration^26^. Notably, Nfix is also important for the correct phenotypical switch and function of MPs upon acute muscle injury. Indeed, Nfix is expressed after ROCK-dependent phagocytosis, inducing a decrease of pro-inflammatory markers and an increase of anti-inflammatory markers^27^.

Here we demonstrated that, unlike what happens in a context of acute damage, Nfix expression by MPs in the dystrophic musculature is associated with the progression of MDs by acting both on myogenic cells and FAPs. Nfix specific deletion in *α-sarcoglycan* null mice (hereafter called *Sgca* null), the murine model for limb-girdle muscular dystrophy 2D, protects myofibers from apoptosis and decreases fibrosis establishment. We showed that MPs deleted for Nfix express more TNFα and less TGFβ1 thus inducing FAPs apoptosis. Moreover, pharmacological inhibition of ROCK in *Sgca* null mice accelerates fibrosis by increasing the number of MPs expressing Nfix. With this study, we demonstrated that Nfix expression by MPs is involved in the progression of MDs *in vivo*. Thus, targeting Nfix in both MPs and myogenic cells could represent a valid approach to delay muscle wasting and fibrosis in MDs.

## Results

### The number of Nfix-positive MPs increases with the progression of Muscular Dystrophy

Since MPs are linked to MDs^11, 20, 28^ and as we demonstrated a pivotal function of Nfix in MPs function during muscle regeneration^27^, we decided to analyze the number of Nfix^+^ MPs during the progression of MD within the *Tibialis Anterior* (TA) and diaphragm (Dia) of two different dystrophic murine models (the *mdx* and the *Sgca* null mice). We observed an increase in the number of MPs expressing Nfix in both TA and Dia muscles of the *mdx* and *Sgca* null mice between 2 and 6 months of life (Figure 1a and b). During muscle regeneration after acute injury, we showed that Nfix is required for Ly6C^-^ anti-inflammatory phenotype acquisition^27^. Although the concept of pro-and anti-inflammatory MPs does not seem to be conserved between acute and chronic muscle injury context, we still decided to look at the number CD64^+^ MPs positive for Nfix within the Ly6C^+^ and Ly6C^-^ population sorted from the TA of *Sgca* null mice. Interestingly, and contrary to what was observed upon acute damage, Nfix was found at comparable levels between Ly6C^+^ and Ly6C^-^ MPs, and its expression increases at the same extent from 2 to 4 months of life in both populations (Figure 1c). These results suggest an involvement of Nfix in MPs in MD progression and confirm that the notion of Ly6C^+^ pro- and Ly6C^-^ anti-inflammatory MPs has different biological meaning in the context of chronic muscle injury contrary to acute muscle injury.

**Fig 1.**
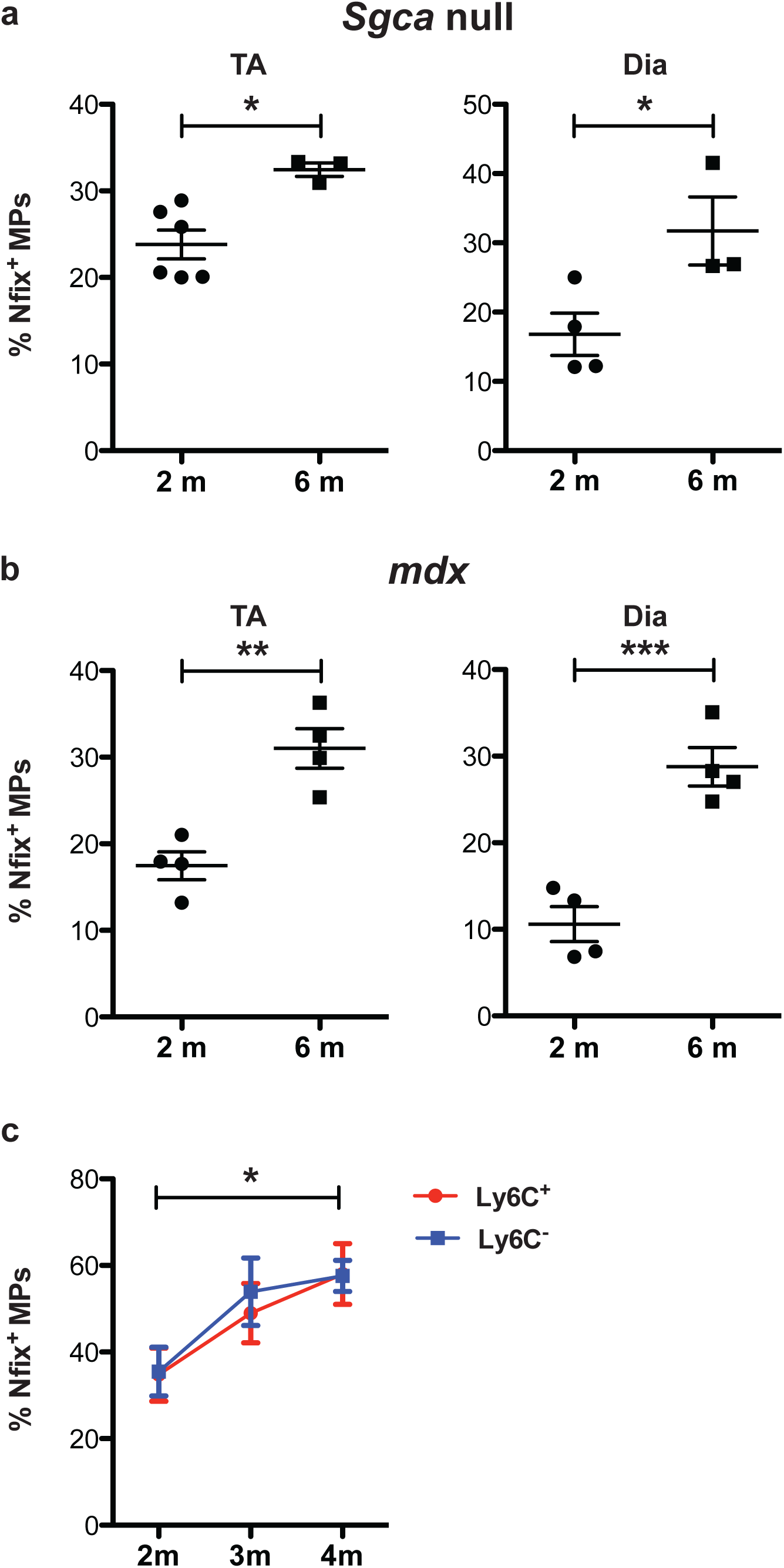
Number of Nfix^+^ MPs increases with age in *Sgca* null and mdx mice. a) Percentage of F4/80^+^ MPs positive for Nfix in *Tibialis Anterior* muscles (TA) and diaphragm (Dia) of *Sgca* null mice at 2 and 6 months of life. b) Percentage of F4/80^+^ MPs positive for Nfix in *Tibialis Anterior* muscles (TA) and diaphragm (Dia) of mdx mice at 2 and 6 months of life. c) Percentage of Ly6C^+^ and Ly6C^-^ sorted MPs positive for Nfix from TA of *Sgca* null mice at 2, 3 and 4 months of life. Statistical significance was determined using a two-tailed Student’s *t*-test, one-way Anova test or two-way Anova test. * p<0,05; ** p<0,01 ; *** p<0,001 ; for c) * p<0,05 Ly6C^-^ 2m vs 4m. Results are means ± SEM of at least three independent experiments.

### Ablation of Nfix in MPs protects the dystrophic muscle

We previously demonstrated that the deletion of Nfix in myogenic cells induces a slower musculature during postnatal muscle development and a delay of skeletal muscle regeneration^24, 25^. Moreover, we observed that silencing of Nfix delays the progression of MD by slowing down muscle regeneration^26^. To understand in which proportion the rescue of dystrophic phenotype observed was due to the lack of Nfix in MPs, we generated *Sgca* null dystrophic mice deleted for Nfix in MPs: the LysM^CRE^:Nfix^fl/fl^:sgca^(-/-)^ mice (hereafter named ΦNfix^(-/-)^:α^(-/-)^). We decided to use the *Sgca* null mice because its phenotype is more severe compared to the *mdx* mice and better recapitulates the progression of DMD patients^4, 29–31^. The LysM^CRE^:Nfix^fl/fl^:sgca^(+/-)^ (hereafter named ΦNfix^(-/-)^:α^(+/-)^) mouse was used as non-dystrophic control and the Nfix^fl/fl^:sgca^(-/-)^ mice (hereafter named Nfix^fl/fl^:α^(-/-)^) as dystrophic control. We first looked at TA histology at 2, 4, and 6 months of life by Hematoxylin and Eosin staining (Figure 2a). Non-dystrophic ΦNfix^(-/-)^:α^(+/-)^ mice displayed a homogenous caliber of the myofibers with myonuclei in peripherical position at all time points analyzed (Figure 2a) which is consistent with our previous observation of a general normal muscle development in ΦNfix^(-/-)^ mice^27^ (Saclier et al 2020). On the contrary, in the dystrophic control Nfix^fl/fl^:α^(-/-)^ mice at 2 months of life, we observed disorganization of the muscle structure characterized by necrotic myofibers, centrally nucleated myofibers (a sign of ongoing regeneration), and areas of inflammatory infiltrates. These features were still observed at 4 months of life with an increased co-presence in the same muscles of small and big fibers. At 6 months of life, an increase of space between myofibers and heterogeneity of myofiber size was present (Figure 2a). At 2 months of life, myofibers in regeneration were detected in the ΦNfix^(-/-)^:α^(-/-)^ dystrophic mice but few inflammatory and necrotic areas were seen. Surprisingly, we did not observe an increase of myofiber size heterogeneity with age and more importantly, there was no appearance of space between myofibers at 6 months of life as observed in the dystrophic control (Figure 2a). We then quantified the caliber of myofibers by measuring the cross-sectional area (CSA). At 2, 4 and 6 months of life, the CSA repartition of the non-dystrophic control exhibited a bell-shape characteristic of WT muscles (Figure 2b). At 2 months of life, the CSA repartition was identical between the Nfix^fl/fl^:α^(-/-)^ and ΦNfix^(-/-)^:α^(-/-)^ mice (Figure 2b). While a shift toward an increase of small myofibers was observed over time in the Nfix^fl/fl^:α^(-/-)^ mice, the CSA repartition of the ΦNfix^(-/-)^:α^(-/-)^ mice remained unmodified up to 6 months of life (Figure 2b). Then, we analyzed the number of regenerative myofibers by quantifying the percentage of centrally nucleated myofibers in the TA at different ages (Figure S1a). Almost no regenerating myofibers were detected in the ΦNfix^(-/-)^:α^(+/-)^ mice. At 2 months of life in the Nfix^fl/fl^:α^(-/-)^ mice, around 86% of myofibers were in regeneration and reached 92% at 6 months of life (Figure S1a). At 2 months, a significant decrease of the percentage of regenerative myofibers (67%) was observed in the ΦNfix^(-/-)^:α^(-/-)^ mice compared to the Nfix^fl/fl^:α^(-/-)^ mice, whereas at 4 and 6 months of life, the number of centrally nucleated myofibers between the two dystrophic murine models was no longer significant. Indeed, the number of regenerative myofibers in the ΦNfix^(-/-)^:α^(-/-)^ mice was around 87% at 6 months of life (Figure S1a). We then analyzed an important hallmark of dystrophies that is myofiber fragility by intraperitoneal (IP) injection of Evans Blue (EB) one day before the sacrifice of the mice. As expected, we did not detect EB^+^ myofibers in the TA of ΦNfix^(-/-)^:α^(+/-)^ mice (Figure 2c and d). In the TA of Nfix^fl/fl^:α^(-/-)^ mice, we observed that damaged myofibers appeared usually in foci, but the quantification revealed a huge heterogeneity in the percentage of necrotic myofibers between animals (from 2 to 18%). This heterogeneity was conserved at 4 months of life, but at 6 months of life few EB^+^ myofibers were present, with the exception of one TA that reached 35% of damaged myofibers (Figure 2d). On the contrary, the percentage of necrotic myofibers in the TA of ΦNfix^(-/-)^:α^(-/-)^ mice was homogenous and strikingly low (no more than 5%) (Figure 2c and d). All these results demonstrate that the deletion of Nfix in MPs has a positive effect on myofiber’s fragility, delaying the appearance of regenerating myofibers and protecting muscle from dystrophic progression.

**Fig 2.**
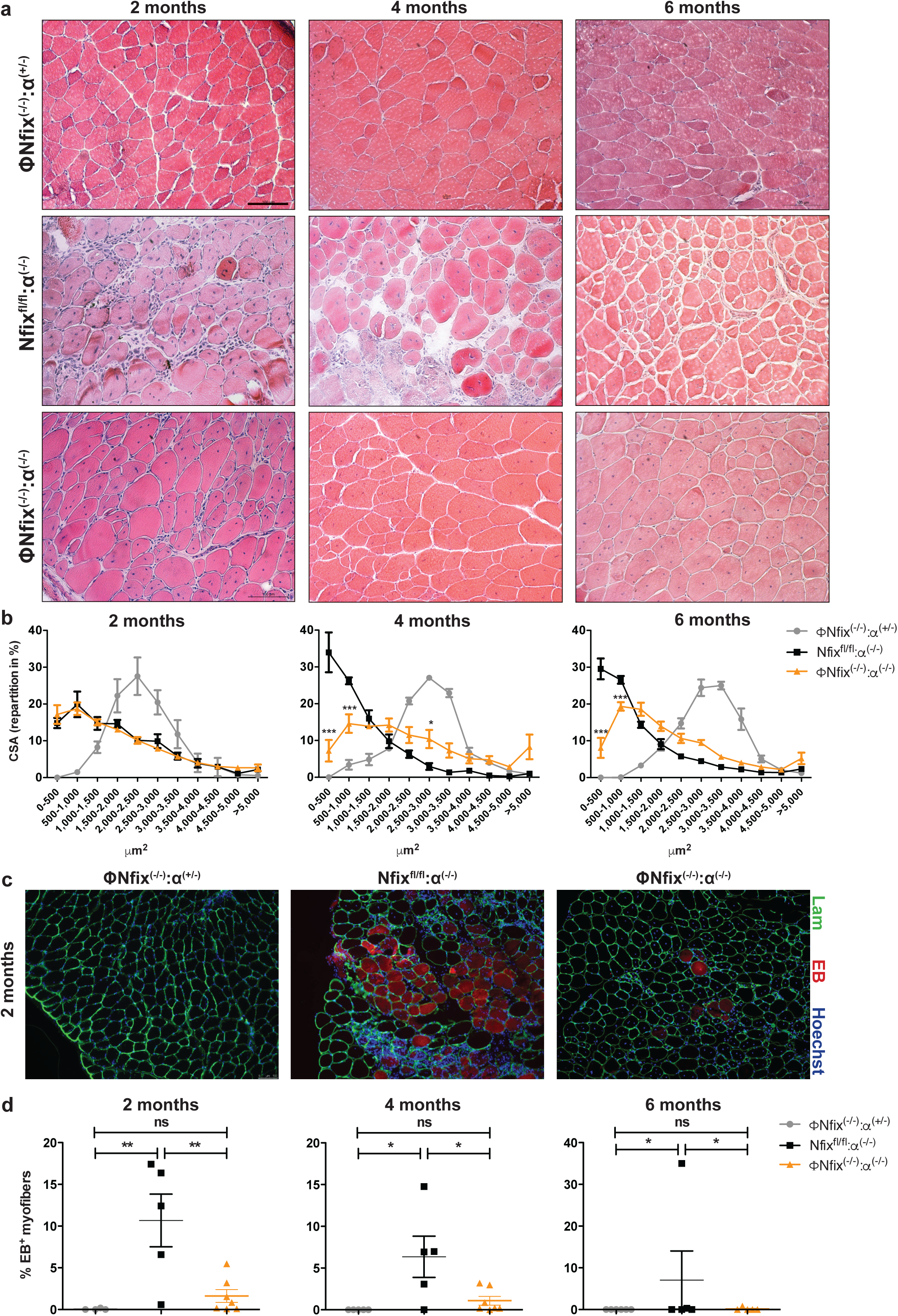
Ablation of Nfix in MPs protects dystrophic muscle. a) Hematoxylin-eosin staining of ΦNfix^(-/-)^:α^(+/-)^, Nfix^fl/fl^:α^(-/-)^ and ΦNfix^(-/-)^:α^(-/-)^ TA muscles at 2, 4 and 6 months of life. b) Repartition in percentage of the Cross-sectional area (CSA). c) Immunostaining for Evan’s blue (EB, red), Lam (green) and Hoechst (blue) of ΦNfix^(-/-)^:α^(+/-)^, Nfix^fl/fl^:α^(-/-)^ and ΦNfix^(-/-)^:α^(-/-)^ TA muscles at 2 months of life. d) Percentage of EB^+^ myofibers at 2, 4 and 6 months of life. EB was injected 24h before sacrifice. Statistical significance was determined using one-way Anova test or two-way Anova test.* p<0,05 ; ** p<0,01 ; for b) * p<0,05 ; *** p<0,001 Nfix^fl/fl^:α^(-/-)^ vs ΦNfix^(-/-)^:α^(-/-)^. Results are means ± SEM of at least three independent experiments. Scale bar = 100μm.

### Fibrosis decreases in Sgca null mice in absence of Nfix in MPs

After cycles of degeneration-regeneration, dystrophic muscle tissue is replaced by fibrotic tissue. We thus performed a Milligan Trichrome staining to highlight ECM in blue in TA of the three mouse models. ECM appeared thin in between the myofibers in TA of the ΦNfix^(-/-)^:α^(+/-)^ mice at 2, 4 and 6 months of life (Figure 3a). At 2 months, the TA of the Nfix^fl/fl^:α^(-/-)^ mice presented large deposits of ECM that increased in size until 6 months (Figure 3a). On the contrary, the TA of ΦNfix^(-/-)^:α^(-/-)^ mice exhibited less ECM than the Nfix^fl/fl^:α^(-/-)^ mice and, importantly no huge areas were observed (Figure 3a). We then performed an immunofluorescence against Collagen I and we quantified the positive area. Around 5% of Collagen I was present in the TA of the ΦNfix^(-/-)^:α^(+/-)^ mice at all time points analyzed (Figure 3b,c). The TA of Nfix^fl/fl^:α^(-/-)^ mice contained more than 20% of collagen I at all the time points (Figure 3c). The TA of ΦNfix^(-/-)^:α^(-/-)^ mice displayed a significant reduction of collagen I deposits compared to the Nfix^fl/fl^:α^(-/-)^ mice over time (Figure 3c). To understand if the improvement of dystrophic muscle histology might also correlate with better muscle functionality, we realized a treadmill test. We did not observe differences in total and weekly time running between Nfix^fl/fl^:α^(-/-)^ and ΦNfix^(-/-)^:α^(-/-)^ mice (Figure S2b), thus suggesting that the lack of Nfix in MPs improves several dystrophic signs of the disease not impinging on muscle performance, which likely depends on muscle cells.

**Fig 3.**
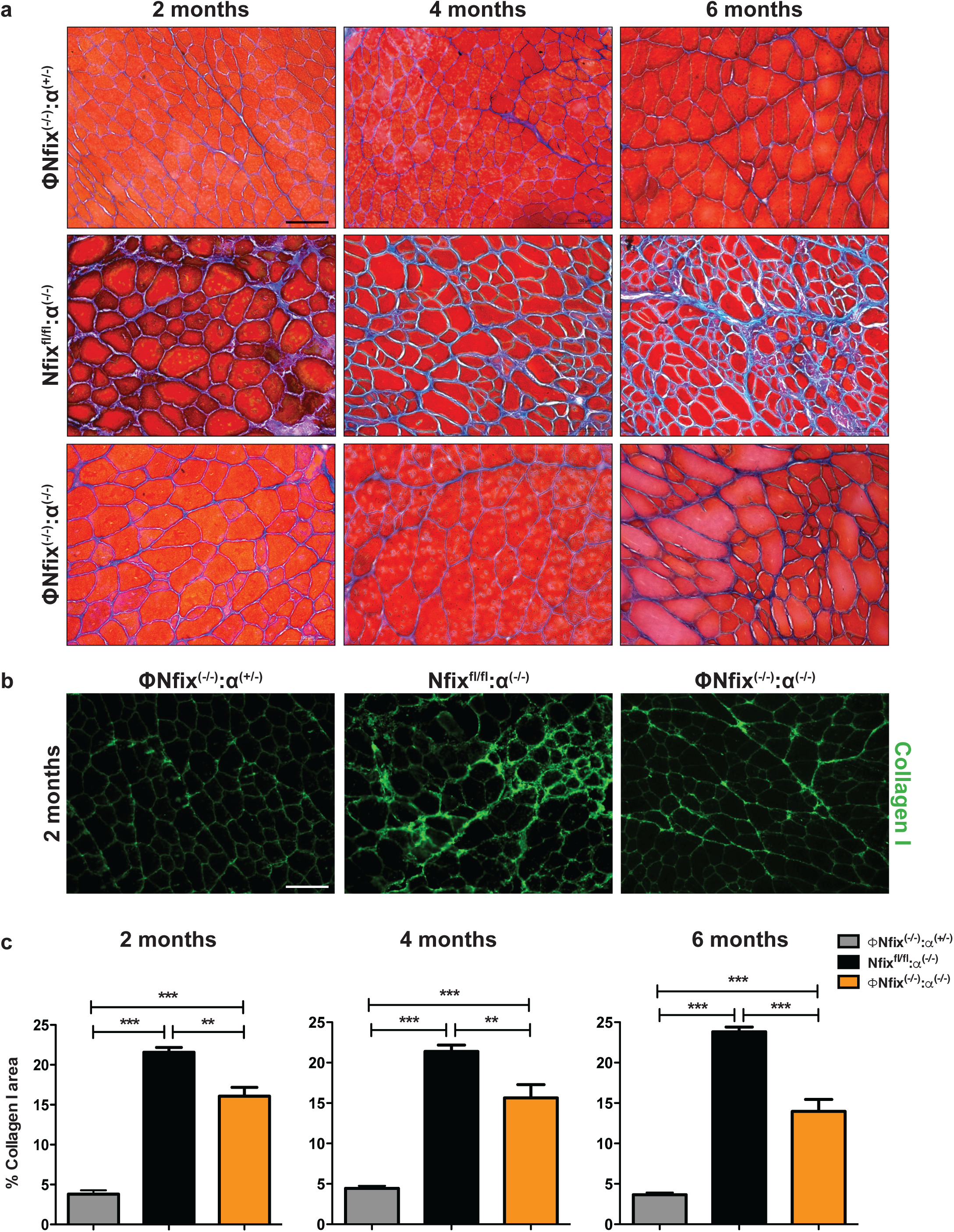
Ablation of Nfix in MPs delays fibrosis. a) Milligan Trichrome staining of ΦNfix^(-/-)^:α^(+/-)^, Nfix^fl/fl^:α^(-/-)^ and ΦNfix^(-/-)^:α^(-/-)^ TA muscles at 2, 4 and 6 months of life. b) Immunostaining for Collagen I (green) of ΦNfix^(-/-)^:α^(+/-)^, Nfix^fl/fl^:α^(-/-)^ and ΦNfix^(-/-)^:α^(-/-)^ TA muscles at 2 months of life. c) Quantification of Collagen I^+^ area at 2, 4 and 6 months of life. Statistical significance was determined using a one-way Anova test.** p<0,01 ; *** p<0,001. Results are means ± SEM of at least three independent experiments. Scale bar = 100μm.

### Ablation of Nfix in MPs preserves diaphragm from fibrosis

We also looked at the Dia histology because this muscle better reflects the dystrophic phenotype^29, 32^. As for the TA, normal muscle histology was observed in the ΦNfix^(-/-)^:α^(+/-)^ mice (Figure 4a). The Dia of Nfix^fl/fl^:α^(-/-)^ mice exhibited a worsen phenotype than TA with an increase of inflammatory cells and space between myofibers, at 2 months of life calcifications were also present (Figure 4a). As for the TA, the Dia of ΦNfix^(-/-)^:α^(-/-)^ mice displayed a better histology compared to the dystrophic control with a decrease of space between myofibers (Figure 4a). After Milligan Trichrome staining, we observed that the Dia of ΦNfix^(-/-)^:α^(+/-)^ displayed thin ECM between myofibers (Figure S1c). As expected, the Dia of the Nfix^fl/fl^:α^(-/-)^ contained large ECM deposits at 2 months, that increased until 6 months of life. At that age, a third of the Dia muscle seemed to be composed by ECM (figure S1c). Compared to the TA, the Dia of ΦNfix^(-/-)^:α^(-/-)^ mice showed some large zones of ECM that were still thinner than those present in the Nfix^fl/fl^:α^(-/-)^ mice (Figure S1c). We then performed immunofluorescence against Collagen deposits and we observed that Collagen I area was higher in both Nfix^fl/fl^:α^(-/-)^ and ΦNfix^(-/-)^:α^(-/-)^ mice compared to the TA (Figure 4b and c versus Figure 3b and c). Notably, with the exception of 2 months, there was a significant decrease of Collagen I area in the Dia of ΦNfix^(-/-)^:α^(-/-)^ mice compared to the Dia of Nfix^fl/fl^:α^(-/-)^ mice (Figure 4b and c). Thus, the ablation of Nfix in MPs also protects Dia from dystrophy.

**Fig 4.**
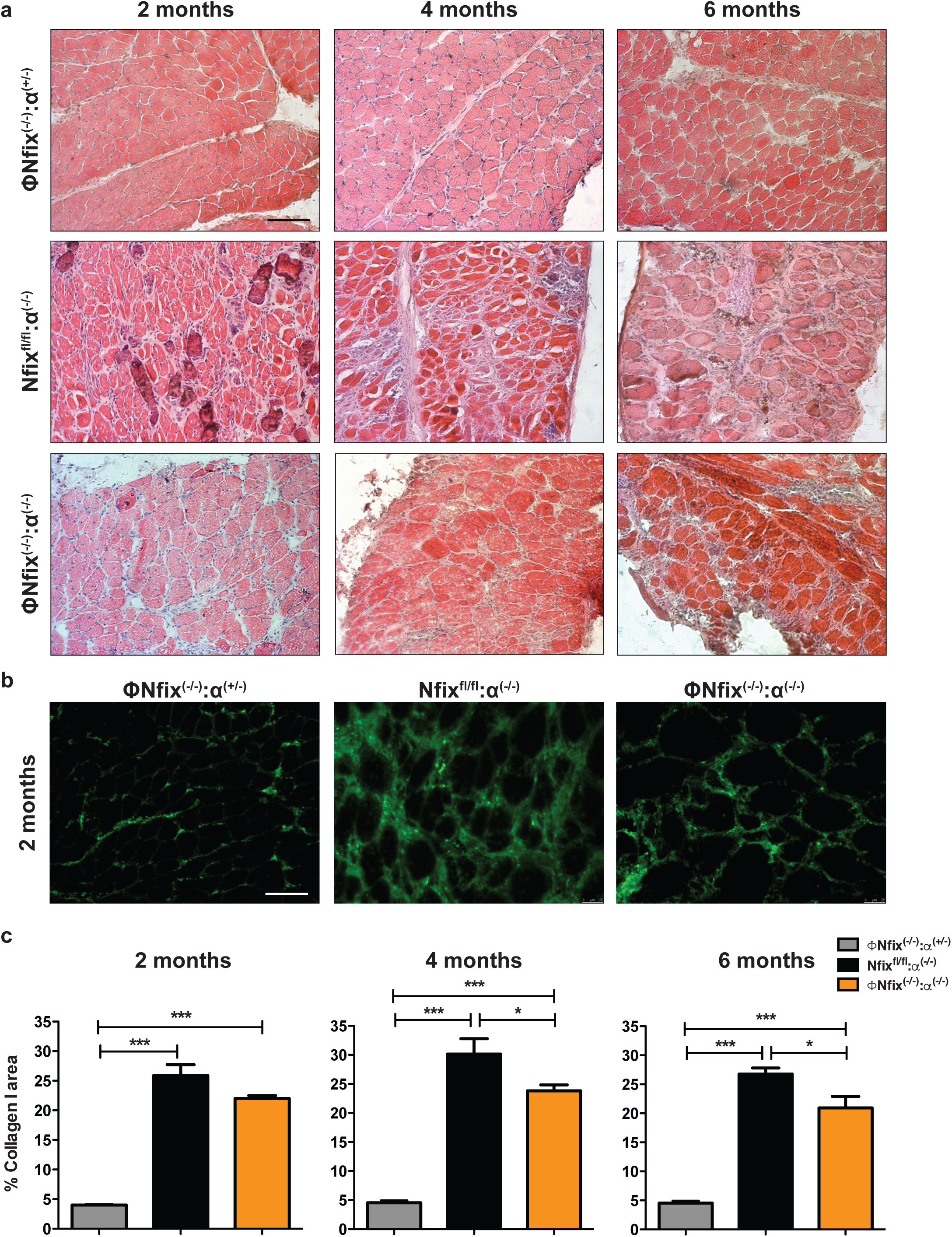
Ablation of Nfix in MPs also preserves Dia from fibrosis. a) Hematoxylin-eosin staining of ΦNfix^(-/-)^:α^(+/-)^, Nfix^fl/fl^:α^(-/-)^ and ΦNfix^(-/-)^:α^(-/-)^ DIA muscles at 2, 4 and 6 months of life. b) Immunostaining for Collagen I (green) of ΦNfix^(-/-)^:α^(+/-)^, Nfix^fl/fl^:α^(-/-)^ and ΦNfix^(-/-)^:α^(-/-)^ Dia muscles at 2 months of life. c) Quantification of Collagen I^+^ area at 2, 4 and 6 months of life. Statistical significance was determined using a one-way Anova test. * p<0,05 ; *** p<0,001. Results are means ± SEM of at least three independent experiments. Scale bar = 100μm.

### Nfix induces pro-fibrotic phenotype of MPs in dystrophic muscle

Within the muscle, FAPs are the main source of ECM. In MDs, MPs induce survival of FAPs thus leading to fibrosis. Interestingly, in the TA of ΦNfix^(-/-)^:α^(-/-)^ mice, we observed a decrease in the number of FAPs (PDGFRα^+^ cells) and MPs (F4/80^+^ cells) compared to Nfix^fl/fl^:α^(-/-)^ mice (Figure 5a). ECM deposits, particularly Collagen I, are due to excessive TGFβ1 signalling, whose expression does not reflect its biological availability. Indeed, TGFβ1 is secreted in a latent form that needs to be cleaved to be active^33^. As seen in other dystrophic mouse models^34, 35^, we have not observed differences in TGFβ1 expression between our two dystrophic models (Figure S2a), but the analysis of its downstream signaling pathway through Smad3 showed a decrease of TGFβ1 pathway activation at 2 and 4 months (Figure 5b). After acute injury, pro-inflammatory MPs express TNFα that induces the apoptosis of FAPs, while TGFβ1 secreted by anti-inflammatory MPs rescue them from TNFα-induced apoptosis and stimulate their proliferation. In a dystrophic context, more than 75% of MPs express TGFβ1 and the consequence of the unbalance between TNFα and TGFβ1 factors leads to FAPs survival^11^. Since we observed a decrease of FAPs number and a decrease of TGFβ1 signaling pathway, we decided to sort FAPs and MPs from hindlimb muscles of Nfix^fl/fl^:α^(-/-)^ and ΦNfix^(-/-)^:α^(-/-)^ mice (Figure 5c, S2b and c). This analysis confirmed the decrease of FAPs and MPs number within muscles of ΦNfix^(-/-)^:α^(-/-)^ mice compared to muscles of Nfix^fl/fl^:α^(-/-)^ mice (Figure S3d). Then we analyzed the number of sorted MPs positive for TNFα or TGFβ1 and the number of sorted FAPs in apoptosis or in proliferation by using aCaspase3 and Ki67 antibodies, respectively (Figure 5d). At 2 and 4 months of life, we observed an increase of MPs positive for TNFα and a decrease of MPs positive for TGFβ1 in ΦNfix^(-/-)^:α^(-/-)^ mice compared to Nfix^fl/fl^:α^(-/-)^ mice (Figure 5e). This modification of cytokines expression did not modify FAPs proliferation but rather induced their apoptosis (Figure 5e). Interestingly, the proportion of Ly6C^+^ and Ly6C^-^ MPs was unchanged between Nfix^fl/fl^:α^(-/-)^ mice and ΦNfix^(-/-)^:α^(-/-)^ mice meaning that the Ly6C receptor was not sufficient in dystrophic context to define pro- and anti-inflammatory MP population (Figure S2e). To conclude, we observed that Nfix is involved in the pro-fibrotic phenotype of MPs and that its deletion rebalances TNFα-TGFβ1 expression promoting apoptosis of FAPs thus delaying fibrosis establishment.

**Fig 5.**
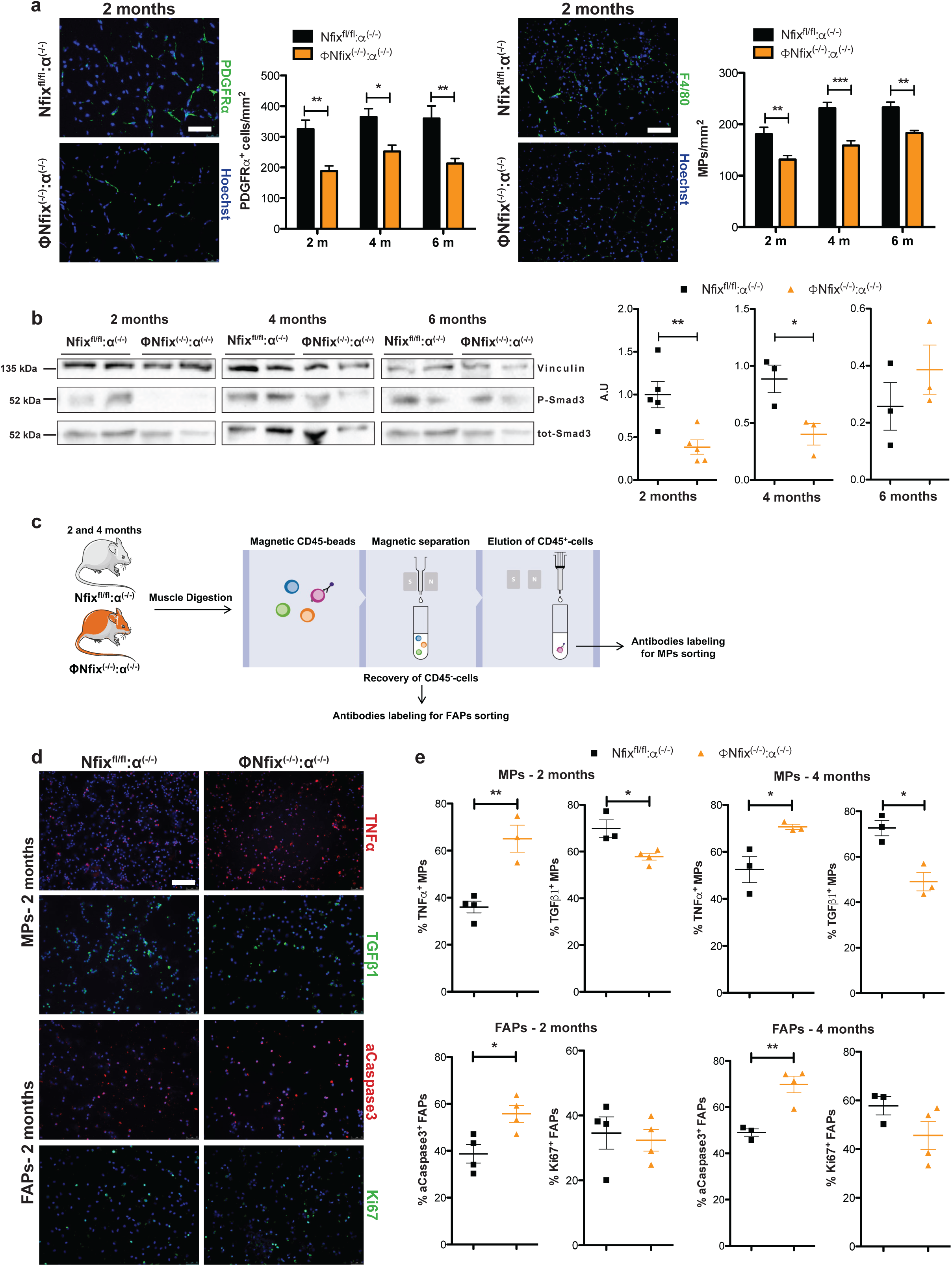
MPs lacking Nfix rescue FAPs apoptosis by reducing TGFβ signalling and expressing TNFα. a) Immunostaining for PDGFRα (green) and Hoechst (blue) of Nfix^fl/fl^:α^(-/-)^ and ΦNfix^(-/-)^:α^(-/-)^ TA muscles at 2 months of life, and number of PDGFRα^+^ cells by mm^2^. Immunostaining for F4/80 (green) and Hoechst (blue) of Nfix^fl/fl^:α^(-/-)^ and ΦNfix^(-/-)^:α^(-/-)^ TA muscles at 2 months of life, and number of F4/80^+^ cells by mm^2^. b) Western Blot of P-Smad3 and tot-Smad3 in Nfix^fl/fl^:α^(-/-)^ and ΦNfix^(-/-)^:α^(-/-)^ TA muscles at 2 months of life and quantification. Western blot were realized in double or membrane were stripped and vinculin was used to normalize. c) Strategy to sort MPs and FAPs by using CD45 magnetic beads on Nfix^fl/fl^:α^(-/-)^ and ΦNfix^(-/-)^:α^(-/-)^ hindlimb muscles at 2 and 4 months of life. d) Immunostaining for TNFα (red) or TGFβ (green) and Hoechst (blue) on cytospined sorted MPs and for aCaspase3 (red) or Ki67 (green) and Hoechst (blue) on cytospined sorted FAPs. e) Percentage of TNFα or TGFβ positive MPs and aCaspase3 or Ki67 positive FAPs sorted from Nfix^fl/fl^:α^(-/-)^ and ΦNfix^(-/-)^:α^(-/-)^ hindlimb muscles at 2 and 4 months of life. Statistical significance was determined using a two-tailed Student’s *t*-test or two-way Anova test. * p<0,05 ; ** p<0,01. Results are means ± SEM of at least three independent experiments. Scale bar = 50 μm for FAPs in a), and 100 μm for MPs in a), and b) and d).

### Pharmacological inhibition of ROCK in dystrophic muscle promotes fibrosis by inducing Nfix in MPs

We previously demonstrated that Nfix is expressed upon RhoA-ROCK dependent phagocytosis in a context of muscle acute injury^27^, thus we asked whether this pathway was conserved in dystrophic musculature and if the stimulation of Nfix expression by MPs could accelerate fibrosis. Therefore, we injected *Sgca* null mice with the ROCK inhibitor Y27632 (then called Y), or its control, 3 times per week for 2 weeks. ΦNfix^(-/-)^:α^(-/-)^ mice were also treated to distinguish the effect of ROCK inhibition on MPs with respect to other cell types present in the muscle (Figure S3a). The injection of Y increased ECM deposits within the TA of *Sgca* null mice compared to control injected *Sgca* null mice, while this effect was not observed on Y injected ΦNfix^(-/-)^:α^(-/-)^ mice compared to control injected ΦNfix^(-/-)^:α^(-/-)^ mice (Figure 6a). In *Sgca* null mice, we observed that Y injection induced an increase of MPs positive for Nfix (Figure 6b). In both control or Y-injected ΦNfix^(-/-)^:α^(-/-)^ mice, around 15% of MPs were found to be positive for Nfix (Figure 6b). This result is in line with our recent study showing that, in an uninjured muscle, resident MPs are positive for Nfix. The percentage of Nfix^+^ resident MPs was around 12% in Nfix^fl/fl^ and ΦNfix^(-/-)^ mice and we did not observe any growth muscle defect in ΦNfix^(-/-)^ mice^27^. While Y injection did not modify the number of MPs/mm^2^ in each murine model, we observed that it increased the number of MPs positive for Nfix only in *Sgca* null mice and not in ΦNfix^(-/-)^:α^(-/-)^ mice, (Figure 6b and S3b). These results mean that the increase of Nfix^+^ MPs in injected *Sgca* null mice was due to a modification of MP phenotype and not to an increase of infiltrated MPs and that this phenomenon is not observed in ΦNfix^(-/-)^:α^(-/-)^ mice. We previously published that the overexpression of Nfix in myofibers exacerbates the dystrophic phenotype and that in embryonic myoblasts, but not in juvenile SCs, ROCK inhibition induces Nfix expression^26, 36^. Here, we observed that the injection of Y in both *Sgca* null mice and ΦNfix^(-/-)^:α^(-/-)^ mice had no effect on Nfix expression by Pax7^+^ SCs but decreased the number of Pax7^+^ SCs/mm^2^ (Figure S3c). On the contrary, we observed an increase in the percentage of Nfix^+^ myofibers suggesting that ROCK inhibition stimulates Nfix expression by myofibers (Figure S3d). While our previous study on the overexpression of Nfix in myofibers showed a decrease of CSA together with an increase of the percentage of centronucleated myofibers, here ROCK inhibition did not modify nor CSA neither the percentage of centronucleated myofibers, or the number of myonuclei/myofibers in control vs Y injected *Sgca* null mice or in control vs Y injected ΦNfix^(-/-)^:α^(-/-)^ mice (Figure S3d). As expected, we observed an increase of CSA in the TA of ΦNfix^(-/-)^:α^(-/-)^ mice compared to *Sgca* null mice and an increase in the percentage of perinucleated myofibers (Figure S4d). Interestingly, Y injection increased Collagen I deposits and the number of FAPs only in *Sgca* null mice but not in ΦNfix^(-/-)^:α^(-/-)^ mice meaning that the fibrotic effect of ROCK inhibition pass through the increase of Nfix in MPs (Figure 6c and d). To conclude, the pharmacological inhibition of ROCK stimulates fibrosis through an increase of Nfix in MPs.

**Fig 6.**
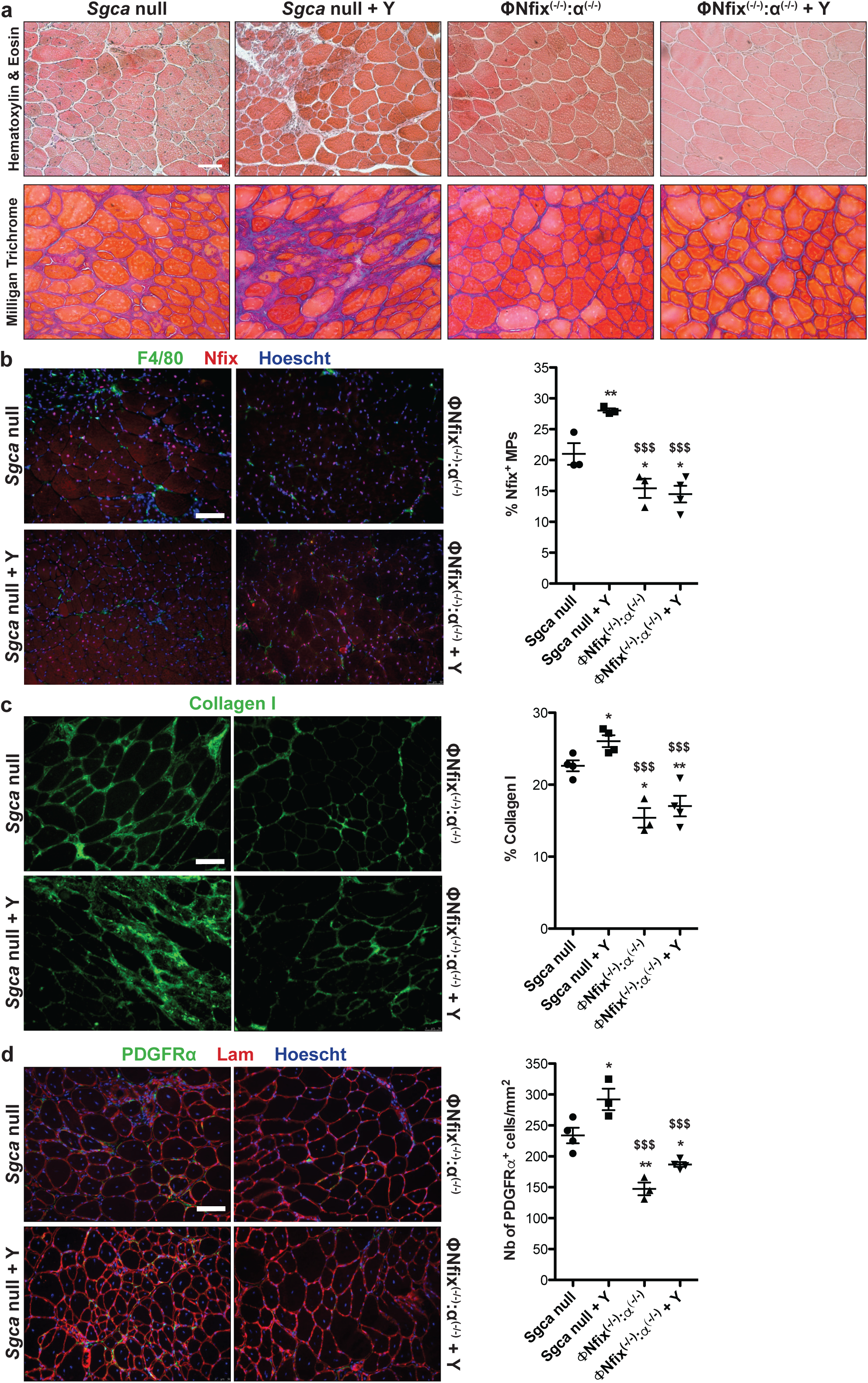
Pharmacological ROCK inhibition induces fibrosis through increase of Nfix^+^ MPs. a) Hematoxylin-eosin and Milligan Trichrome staining of *Sgca* null and ΦNfix^(-/-)^:α^(-/-)^ TA muscles i.p. injected with physiological water or Y27632 (called Y). b) Immunostaining for Nfix (red), F4/80 (green) and Hoechst (blue) of *Sgca* null and ΦNfix^(-/-)^:α^(-/-)^ TA muscles i.p. injected with physiological water or Y, and percentage of double Nfix^+^/F4/80^+^ cells. c) Immunostraining for Collagen I (green) of *Sgca* null and ΦNfix^(-/-)^:α^(-/-)^ TA muscles i.p. injected with physiological water or Y, and percentage of Collagen I^+^ area. d) Immunostaining for Lam (red), PDGFRα (green) and Hoechst (blue) of *Sgca* null and ΦNfix^(-/-)^:α^(-/-)^ TA muscles i.p. injected with physiological water or Y, and number of PDGFRα^+^ cells by mm^2^. Statistical significance was determined using a two-tailed Student’s *t*-test or one-way Anova test. * vs *Sgca* null: * p<0,05 ; ** p<0,01. $ vs *Sgca* null +Y: $$ p<0,01 ; $$$ p<0,001. Results are means ± SEM of at least three independent experiments. Scale bar = 50 μm.

## Discussion

MDs are incurable diseases and, although several approaches such as cellular or gene therapies aim to treat MDs, these demonstrated to be of low clinical benefit so far^37–39^. While most of these strategies are obviously focused on muscle cells, which bear the genetic defect, it is now clear in the field that we should also take into account the cellular environment. Indeed, several studies demonstrated the importance of non-muscle cells in the progression of MDs, particularly MPs^28, 40–43^. In this study, we demonstrated that the expression of transcription factor Nfix induces pro-fibrotic features in MPs in a context of chronic muscle injury. Indeed, MPs deleted for Nfix secrete more TNFα and less TGFβ1 pushing FAPs toward apoptosis. In addition, pharmacological treatment of *Sgca* null mice with ROCK inhibitor shapes MPs toward Nfix expression inducing FAPs persistence and exacerbating fibrosis.

The genetic deletion of Nfix in MPs rescues dystrophic muscle by decreasing myofibers death, indeed some studies already demonstrated the protective capacity of MPs on extensive tissue damage or cellular death^44–46^. Thus, in our ΦNfix^(-/-)^:α^(-/-)^ murine model, myofibers probably enter lately in degeneration/regeneration cycles explaining the decrease of regenerative centronucleated myofibers at 2 months of life compared to Nfix^fl/fl^:α^(-/-)^ mice. This protective effect also acts on myofibers caliber: while an increase of the percentage of small caliber was observed in Nfix^fl/fl^:α^(-/-)^ mice in time, the repartition of myofiber size was unchanged from 2 to 6 months of life in ΦNfix^(-/-)^:α^(-/-)^ mice. The mechanisms linking MPs to fibrosis start to be better known now. In particular, it has been seen that MPs are the main source of TGFβ1 that promotes FAPs survival and Collagen I expression, but also reprograms endothelial cells and SCs toward a fibrotic phenotype^10, 11, 20, 47^. Here we observed that Nfix is a profibrotic factor in MPs since its absence rebalances cytokines secretion toward more TNFα and less TGFβ1. Importantly, we did not observe a difference in terms of Ly6C^+^ and Ly6C^-^ MP populations in our two murine models, consolidating the general idea in the field that the phenotypical classification as pro- and anti-inflammatory populations based on Ly6C marker is not relevant in the case of chronic muscle injury^11, 20^. Moreover, since asynchronous muscle regeneration has been associated with inflammation and fibrotic signatured^48^, our study shows that Nfix deletion in MPs rescues MDs from fibrosis directly by inducing FAP apoptosis and indirectly by protecting myofibers from injury.

We previously demonstrated that RhoA/ROCK-dependent phagocytosis pathway induces Nfix expression during skeletal muscle regeneration upon acute injury^27^. Here, we observed that pushing Nfix expression in MPs by treating *Sgca* null mice with ROCK inhibitor accelerates fibrosis accumulation and establishment. Modulation of ROCK pathway is of high interest in cancer or in hypertension since its inhibition decreases cell migration but also induces vasodilatation^49, 50^. RhoA/ROCK pathway also plays a role in myogenic cells’ fate. In C2C12 cell line, RhoA activity is high during proliferation, then decreases during the first phase of differentiation and increases at a later stage during myofibers growth^51^. In rat, RhoA overexpression keeps SCs in myogenic fate and stimulates myogenin expression^52^. Recently, two studies demonstrated that RhoA is important for niche retention and maintenance of the myogenic capacity of SCs. Indeed, inhibition of ROCK accelerates muscle regeneration after acute injury^53, 54^. In overload conditions, RhoA signaling in myofibers is required for proper myofibers hypertrophy. Particularly, RhoA loss decreases the number of myonuclei/myofibers by impairing the fusion of SCs with myofibers^55^. In our two murine models, ROCK inhibition does not modify the number of SCs expressing Nfix but we observed a decrease in the number of SCs/mm^2^ that is consistent with previous studies^53, 54^. On the contrary, we observed an increase of Nfix^+^ myofibers suggesting that the ROCK/Nfix pathway is conserved in muscle fibers. While overexpression of Nfix in myofibers exacerbates dystrophy by decreasing myofibers caliber and increasing the percentage of centronucleated myofibers^26^, the increase of Nfix^+^ myofibers in both Y injected *Sgca* null mice and ΦNfix^(-/-)^:α^(-/-)^ mice has no effect on these two parameters. Similarly, we did not observe a modification of myonuclei number within myofibers. Thus, the effect of ROCK inhibition on SCs number and percentage of Nfix expressing myofibers does not correlate with a modification of muscle features. This could be explained by the fact that chronic muscle injury is a completely different context compared to overload-induced hypertrophy or muscle regeneration after acute injury. To note, since we previously observed that the exacerbation of the dystrophic phenotype in the dystrophic mice overexpressing Nfix in myofibers correlates with an increase of Nfix levels in muscle cells^26^, it is also possible that our treatment was not sufficient to reach a level of Nfix that might be deleterious for dystrophic muscle.

Dystrophin/utrophin double knockout mouse model treated with ROCK inhibitor demonstrate a reduction of heterotopic ossification in skeletal muscle and heart^56, 57^. Contrary to the *mdx* and *Sgca* null mice, the dystrophin/utrophin double knockout mouse model has a more severe phenotype which is closer to the one of DMD patients^58–60^ (Janssen P M 2005, Deconinck cell 1997, Grady cell 1997). Particularly, they develop calcification within skeletal muscle^61^, while in *Sgca* null mice calcification represents less than 1% of gastrocnemius, quadriceps, triceps, or DIA area^29^. The formation of heterotopic ossification seems to be linked to the total absence of dystrophin expression^62^, indeed in *Sgca* null mice, α-sarcoglycan is absent but, even if reduced, dystrophin is still present in dystrophic muscles^30^. This could explain differences observed upon ROCK inhibition treatment between these two murine models. Nevertheless, we observed that the increase of ECM deposits and number of FAPs occur only in Y treated *Sgca* null mice and not in ΦNfix^(-/-)^:α^(-/-)^ mice and, that these increases correlate with the number of MPs expressing Nfix without modifying the total number of MPs present in the muscle. This demonstrates that modifying MP phenotype through Nfix expression is sufficient to promote fibrosis. Therapeutically speaking, to delay the progression of MDs it seems preferable to modulate MP phenotype rather than reduce their number within the muscle. Indeed, the decrease of MP infiltration through to the deletion of CCR2 in *mdx* has beneficial effects in terms of necrotic myofibers and fibrosis reduction in the short-term (12 weeks) which are lost at 6 months of life^13, 63^. Here, we observed an improvement of dystrophic histopathology that is conserved until 6 months of life. Importantly, the deletion of Nfix in myogenic cells leads to benefits in terms of oxidative phenotype and improvement of muscle function^26^, while deletion of Nfix in MPs plays more on the protection of muscle fibers and inhibition of fibrosis. Therefore, targeting Nfix in both myogenic cells and MPs could be useful to preserve musculature in MD patients. Interestingly, several technics using nanoparticles, liposomes or nanogels to encapsulate drugs and use MPs as drug carriers, but also as drug targets, are now under development^64–66^. Moreover, it has been recently demonstrated that MPs efficiently uptake large amounts of phosphorodiamidate morpholino oligomers to deliver them to myoblasts in dystrophic lesions^67^. Thus, we can speculate that upon Nfix inhibition, MPs would change phenotype and protect dystrophic muscle, even by delivering drugs to myogenic cells. Of course, the question of the muscle specificity of drugs that selectively inhibit Nfix in both cell types has to be addressed, but it could be a promising way to delay both muscle loss and fibrosis progression in MDs.

## Materials and Methods

### Animal models and in vivo experimentations

*Mdx*, *Sgca* null, Nfix^fl/fl^:α^(-/-)^ (for Nfix^fl/fl^:sgca^(-/-)^), ΦNfix^(-/-)^:α^(+/-)^ (for LysM^CRE^:Nfix^fl/fl^:sgca^(+/-)^) and ΦNfix^(-/-)^:α^(-/-)^ (for LysM^CRE^:Nfix^fl/fl^:sgca^(-/-)^) mice were used in this study. ΦNfix^(-/-)^:α^(-/-)^ (for LysM^CRE^:Nfix^fl/fl^:sgca^(-/-)^) mice were generated crossing ΦNfix^(-/-)^ (or LysM^CRE^:Nfix^fl/fl^) mice with the *Sgca* null mice^27, 30^. All mice having the LysM^CRE^ gene were heterozygous for it. For some experiments, 10 weeks-old *Sgca* null mice and ΦNfix^(-/-)^:α^(-/-)^ (ofr LysM^CRE^:Nfix^fl/fl^:sgca^(-/-)^) mice were injected IP with physiological water or Y-27632 ROCK1 inhibitor (10mg/kg, Santacruz sc-281642) 3 times per week for 2 weeks. Mice were kept in pathogen-free conditions and all procedures were conformed to Italian law (D. Lgs n° 2014/26, implementation of the 2010/63/UE) and approved by the University of Milan Animal Welfare Body and by the Italian Minister of Health.

### Histology and immunofluorescence analyses

Fascia of TA muscles was removed and muscles frozen in liquid nitrogen-cooled isopentane (VWR) and placed at −80°C until cut. 8 μm-thick cryosections were stained for Hematoxylin and Eosin, Milligan’s Trichrome or used for immunofluorescence.

Hematoxylin and Eosin staining was processed according to standard protocols. For Milligan’s trichrome staining, sections were fixed for 1 h with Bouin’s fixative (Sigma-Aldrich) and rinsed for 1 hr under running water. Sections were then dehydrated to 95% EtOH in graded ethanol solutions, successively passed in 3% potassium dichromate (Sigma-Aldrich) for 5 min, washed in distilled water, stained with 0.1% acid fuchsin (Sigma-Aldrich) for 30 s, washed again in distilled water, passed in 1% phosphomolybdic acid (Sigma-Aldrich) for 3 min, stained with Orange G (2% in 1% phosphomolybdic acid; Sigma-Aldrich) for 5 min, rinsed in distilled water, passed in 1% acetic acid (VWR) for 2 min, stained with 1% Fast Green for 5 min (Sigma-Aldrich), passed in 1% acetic acid for 3 min, rapidly dehydrated to 100% EtOH, and passed in xylene before mounting with Eukitt (Bio-Optica).

For immunofluorescence analysis, sections or cells were fixed for 15 min with 4% paraformaldehyde (except for F4/80). Then samples were permeabilized with 0.5% Triton X-100 (Sigma-Aldrich) in PBS for 10 min and blocked with 4% BSA (Sigma-Aldrich) in PBS at RT for 1 hr. Primary antibodies were incubated O/N at 4°C in PBS. After three washes of 5 min with PBS, samples were incubated with secondary antibodies (1:500, Jackson Laboratory. Fluorochromes used: 488 and 594) and Hoechst (1:500, Sigma-Aldrich) in PBS for 45 min at RT, then washed four times for 5 min with PBS and mounted with Fluorescence Mounting Medium (Dako). For Nfix-F4/80 double immunolabeling, cryosections were labelled with antibodies against F4/80 (1:400, Novus Biologicals NB300-605) overnight at 4°C and Nfix labelling using (1:200, Novus Biologicals NBP2-15038) antibody was performed for 2 hr at 37°C. Pax7 immunolabelling was done on fresh cut muscle sections and antigen retrieval was performed by incubating muscle sections in boiling 10 mM citrate buffer pH6 for 20 min. Muscle sections were then incubated O/N with Pax7 (1:2, Hybridome). Anti-collagen I (1:200, Southern Biotech 1310-01), anti-PDGFRα (1/500, R&D Systems AF1062) and anti-Laminin (1:300, Sigma-Aldrich L9393) antibodies were used on muscle sections. Anti-TNFα (1:100, Genetex GTX54419) anti-TGFβ1 (1:100, abcam ab64715), anti-aCaspase3 (1:300, Cell Signaling 9661S) and anti-Ki67 (1:50, BD Biosciences 550609) antibodies were used on FAPs or MPs sorted cells.

### Evans Blue Dye uptake measurement

Evans Blue Dye (Sigma-Aldrich, solution at 5 mg/ml) was injected 24 hr before the sacrifice in intraperitoneal (10uL per g of mouse). Positivity for Evan’s blue dye was revealed through its autofluorescence. Sections were fixed with acetone (VWR) for 10 min at −20 °C, permeabilized in 0.5% Triton X-100 (Sigma-Aldrich) for 10 min, saturated 1 hr in 4% BSA and incubated O/N with rabbit anti-Laminin antibody (1:300, Sigma-Aldrich) to reveal myofiber outlines. The day after, sections were washed, incubated with a goat anti-rabbit 488 secondary antibody (1:250, Laboratory) and Hoechst (1:500, Sigma-Aldrich) in PBS for 45 min at RT, then washed four times for 5 min with PBS and mounted with Fluorescence Mounting Medium (Dako). Measurement of the percentage of Evans Blue Dye uptake was performed counting the number of Evan’s blue dye positive fibers on total muscle section reconstructions, using Image J software.

### RNA extraction and qRT-PCR

RNA was isolated from TA by using TRIzol Reagent (Invitrogen 15596026) according to the manufacturer’s instructions. RNA was quantified using a NanoPhotoneter (Implen). For retro-transcription, 500 ng of RNA was used with the iScript Reverse Transcription Supermix for RT-quantitative qPCR (Bio-Rad 1708840). For qRT PCR, cDNA was diluted 1:10 and 5 μl of the diluted cDNA was loaded in a total volume of 20 μl (SYBR Green Supermix (Bio-Rad 172-5124) and run on the Bio-RAD CFX Connect Real-Time System. The relative quantification of gene expression was determined by comparative CT method, and normalized to ACTB. Primers used are:

TGFβ1 for AACAACGCCATCTATGAGAAAACC;

TGFβ1 rev CCGAATGTCTGACGTATTGAAGAA;

ACTB for GTAAAACGCAGCTCAGTAACAGTCCG;

ACTB rev CTCTGGCTCCTAGCACCATGAAGA.

### Protein extraction and Western Blot

Protein extracts were obtained from TA lysed using RIPA buffer (10 mM Tris-HCl pH 8.0, 1 mM EDTA, 1% Triton-X, 0.1% sodium deoxycholate, 0.1% sodium dodecylsulphate (SDS), 150 mM NaCl, in deionised water) plus protease and phosphatase inhibitors for 30 min on ice. Then, samples were centrifuged at 11,000 g for 10 min at 4°C, and the supernatants collected for protein quantification (DC Protein Assays Bio-Rad 5000111). 40 μg protein of each sample was denaturated at 95°C for 5 min using SDS PAGE sample-loading buffer (100 mM Tris pH 6.8, 4% SDS, 0.2% bromophenol blue, 20% glycerol, 10 mM dithiothreitol) and loaded into 8% SDS acrylamide gels. After electrophoresis, the protein was blotted into nitrocellulose membranes (Protran nitrocellulose transfer membrane; Whatman) for 2 hr at 70 V at 4°C. Membrane were then blocked for 1 hr with 5% milk in Tris-buffered saline plus 0.02% Tween20 (Sigma-Aldrich). Membranes were incubated with the primary antibodies O/N at 4°C, using the following antibodies: rabbit anti-P-Smad3 (1:2000, abcam ab52903), rabbit anti-Tot-Smad3 (1:1000, Cell Signaling 9523) and mouse anti-vinculin (1:2500, Sigma-Aldrich V9131). After incubation with the primary antibodies, the membranes were washed 3 times for 5 min and incubated with the secondary antibodies (1:10000, IgG-HRP, Bio-Rad) for 45 min at RT, and then washed again 5 times for 5 min. Bands were revealed using ECL detection reagent (ThermoFisher), with images acquired using the ChemiDoc MP system (Bio-Rad). After incubation with rabbit anti-P-Smad3 antibody, the 4 months TA sample membrane was stripped with mild stripping solution (200 mM glycine, 1% SDS, 10% Tween 20, pH = 2,2). After verification of correct stripping by incubating membrane with anti-rabbit-HRP secondary antibody for 45min at RT, membrane was then incubated with anti-Tot-Smad3 antibody O/N. The Image Lab software was used to measure and quantify the bands of at least three independent samples. The obtained absolute quantity was compared with the reference band (Vinculin) and expressed in the graphs as normalised volume (Norm. Vol. Int.).

### Isolation of MPs and FAPs from skeletal muscle

Fascia of the TA muscles was removed. Muscles were dissociated and digested in RPMI medium containing 0,2% of collagenase B (Roche Diagnostics GmbH 11088815001) at 37°C for 1 hr and passed through a 70 μm and a 30 μm cell strainer (Miltenyi Biotec). CD45^+^ cells were isolated using magnetic beads (Miltenyi Biotec 130-052-301) and incubated with FcR blocking reagent (Miltenyi Biotec 130-059-901) for 20 min at 4°C in PBS 2% FBS. Then CD45^+^ cells were then incubated with Ly6C-PE (eBioscience 12-5932) and CD64-APC (BD Pharmingen 558539) antibodies for 30 min at 4°C. CD45^-^ were incubated with CD31-FITC (1:1000, Invitrogen 11-0311-82), CD45-FITC (1:1000, Invitrogen 11-0451-82), ITAG7(a7)-APC (1:1000, Invitrogen MA5-23555) and Sca1-PeCy7 (1:5000, Invitrogen 25-5981-82) antibodies for 30min at 4°C. MPs (CD64^+^ cells) and FAPs (CD31^-^/CD45^-^/a7^-^/Sca1^+^ cells) were analysed and sorted using a FACS Aria III cell sorter (BD Biosciences) (gating strategy is shown Figure S3). In some experiments, Ly6C^+^ and Ly6C^-^ MPs, or CD64^+^ MPs and FAPs were cytospined on starfrost (Knitterglaser) slides and immunostained.

### Treadmill test

For Treadmill test functional assay, 4-week-old mice were exercised three times, once a week, before recording of their performances. Treadmill test was therefore performed starting from 7-week-old mice, once a week for 6 weeks. The test was conducted on a treadmill (Bioseb) with a 10% incline, starting from a speed of 6 cm/s and increasing it by 2 cm/s every 2 min. For each test, the time to exhaustion of each mouse was measured.

### Image acquisition and quantification

Images were acquired with an inverted microscope (Leica-DMI6000B) equipped with Leica DFC365FX and DFC400 cameras and 10X, 20X and 40X magnification objectives. For each condition of each experiment, at least 8 fields chosen randomly were counted. Measurement of central nucleation and myofiber CSA was performed using Image J software. At least 500 myofibers were analysed. Collagen I quantification was performed using a Macro in ImageJ to identify and quantify positive areas. The number of labelled MPs or FAPs was calculated using the cell tracker in ImageJ software and expressed as a percentage of total MPs or FAPs.

### Statistical Analysis

All data shown in graph are expressed as mean ± SEM. All experiments were performed using at least three different animals in independent experiments. Statistical analysis was performed using two-tailed unpaired Student’s t-Test, one-way ANOVA or two-way ANOVA. *P<0,05; **P<0,01; ***P<0,001; $P<0,05; $$P<0,01; $$$ P<0,001; confidence intervals 95%, alpha level 0,05.

## Acknowledgements

We thank R. Gronostajski for the kind exchange of information and animal models. We are also grateful to B. Chazaud and R. Mounier for helpful discussions and the exchange of animal models. We thank G. Angelini and G. Mura for carefully reading the manuscript. This research was funded by the European Community, ERCStG2011 (RegeneratioNfix 280611) and the Association Française contre les Myopathies AFM-Telethon (Grant number 20002).

## Author contributions

M.S. designed and performed experiments, analysed data, interpreted results and wrote the manuscript. G.T. and C.B. performed experiments and analysed data. G.M. supervised the work and wrote the paper.

---

**Fig S1.**
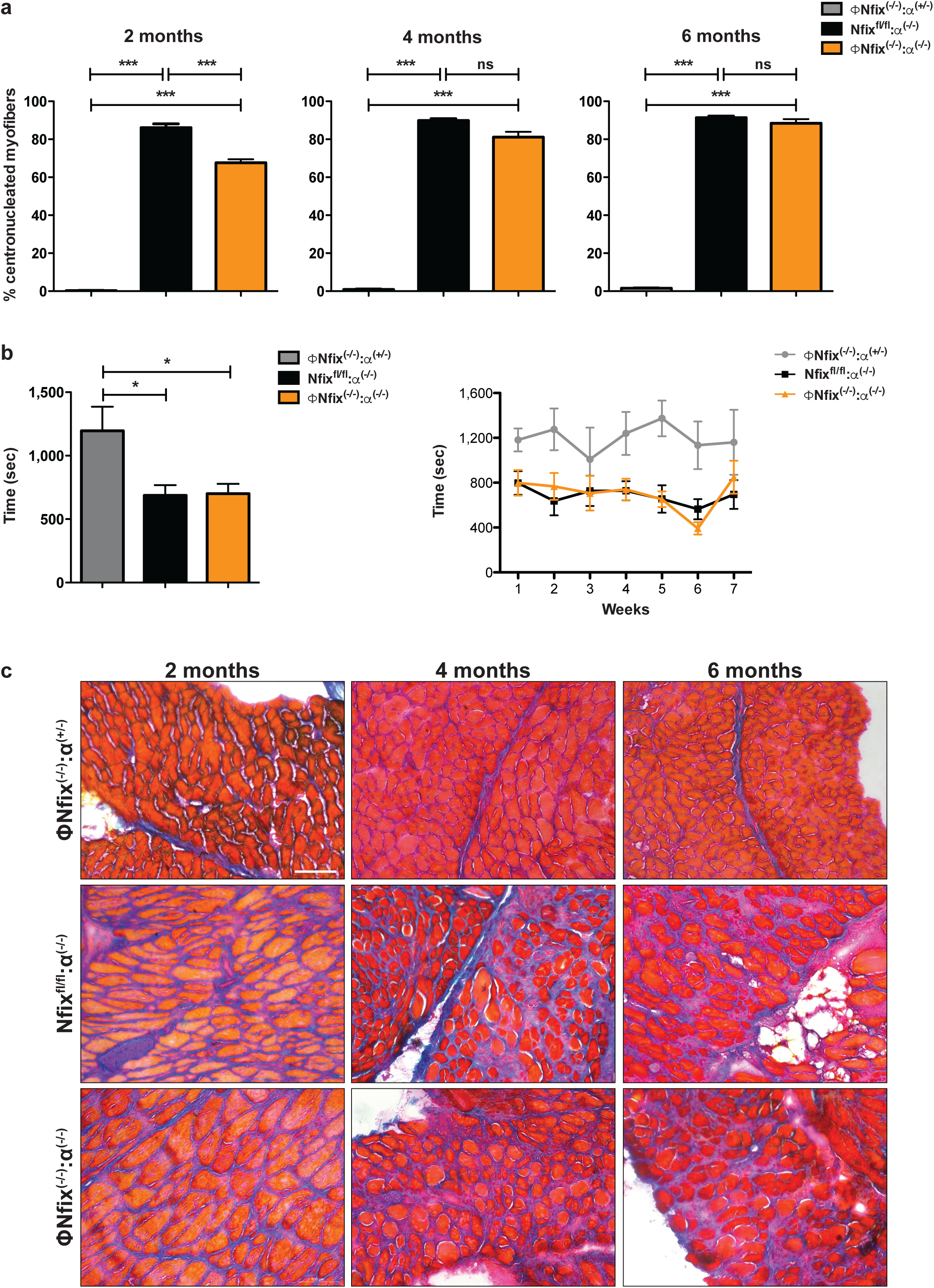
Ablation of Nfix in MPs protects dystrophic muscle but does not improve running performance. a) Percentage of centronucleated myofibers of ΦNfix^(-/-)^:α^(+/-)^, Nfix^fl/fl^:α^(-/-)^ and ΦNfix^(-/-)^:α^(-/-)^ TA muscles at 2, 4 and 6 months of life. b) Treadmill test totality of the measurements and over time on ΦNfix^(-/-)^:α^(+/-)^, Nfix^fl/fl^:α^(-/-)^ and ΦNfix^(-/-)^:α^(-/-)^ mice. c) Milligan Trichrome staining of ΦNfix^(-/-)^:α^(+/-)^, Nfix^fl/fl^:α^(-/-)^ and ΦNfix^(-/-)^:α^(-/-)^ Dia muscles at 2, 4 and 6 months of life. Statistical significance was determined using one-way Anova test or two-way Anova test.* p<0,05 ; *** p<0,001. Results are means ± SEM of at least three independent experiments. Scale bar = 100μm.

**Fig S2.**
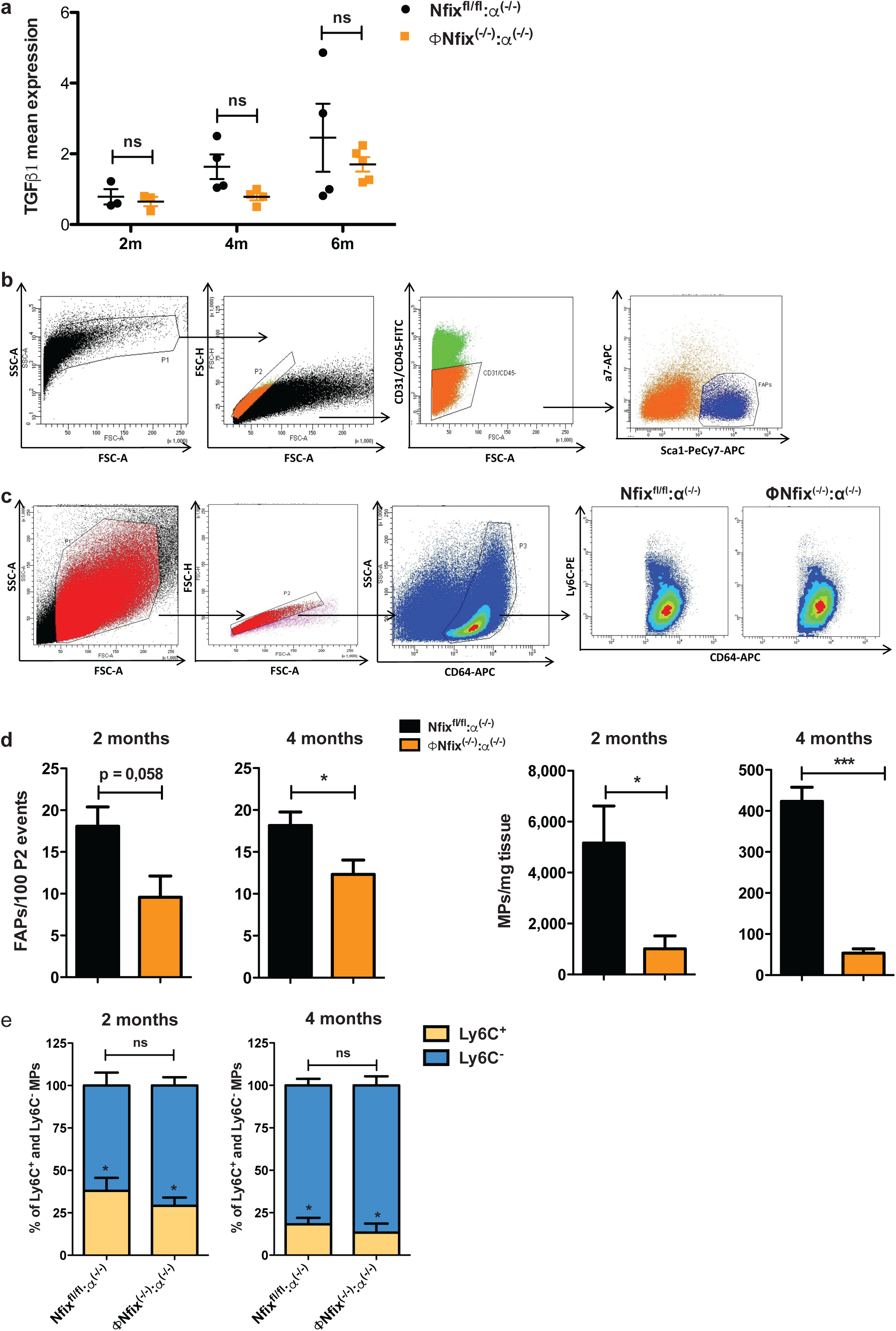
TGFβ1 expression, gating strategy and MPs and FAPs number. a) TGFβ1 expression by RT-qPCR of Nfix^fl/fl^:α^(-/-)^ and ΦNfix^(-/-)^:α^(-/-)^ TA muscles at 2, 4 and 6 months of life. b) Nfix^fl/fl^:α^(-/-)^ and ΦNfix^(-/-)^:α^(-/-)^ TA muscles were digested using Collagenase B and CD45^-^ cells were selected with magnetic beads. CD45^-^ cells were incubated with CD31-FITC, CD45-FITC, ITAG7(a7)-APC and Sca1-PeCy7 antibodies. FAPs are CD31^-^/CD45^-^/a7^-^/Sca1^+^. c) Nfix^fl/fl^:α^(-/-)^ and ΦNfix^(-/-)^:α^(-/-)^ TA muscles were digested using Collagenase B and CD45^+^ cells were selected with magnetic beads. CD45^+^ cells were incubated with CD64-APC and Ly6C-PE. MPs are CD64^+^ cells. Representative FACS gate of Ly6C^+^ and Ly6C^-^ MPs from Nfix^fl/fl^:α^(-/-)^ and ΦNfix^(-/-)^:α^(-/-)^ TA muscles at 2 months of life. d) Number of FAPs and MPs in Nfix^fl/fl^:α^(-/-)^ and ΦNfix^(-/-)^:α^(-/-)^ TA muscles at 2 and 4 months of life. e) Percentage of Ly6C^+^ and Ly6C^-^ MPs in Nfix^fl/fl^:α^(-/-)^ and ΦNfix^(-/-)^:α^(-/-)^ TA muscles at 2 and 4 months of life. Statistical significance was determined using a two-tailed Student’s *t*-test, one-way Anova test or two-way Anova test.* p<0,05 ; *** p<0,001. Results are means ± SEM of at least three independent experiments.

**Fig S3.**
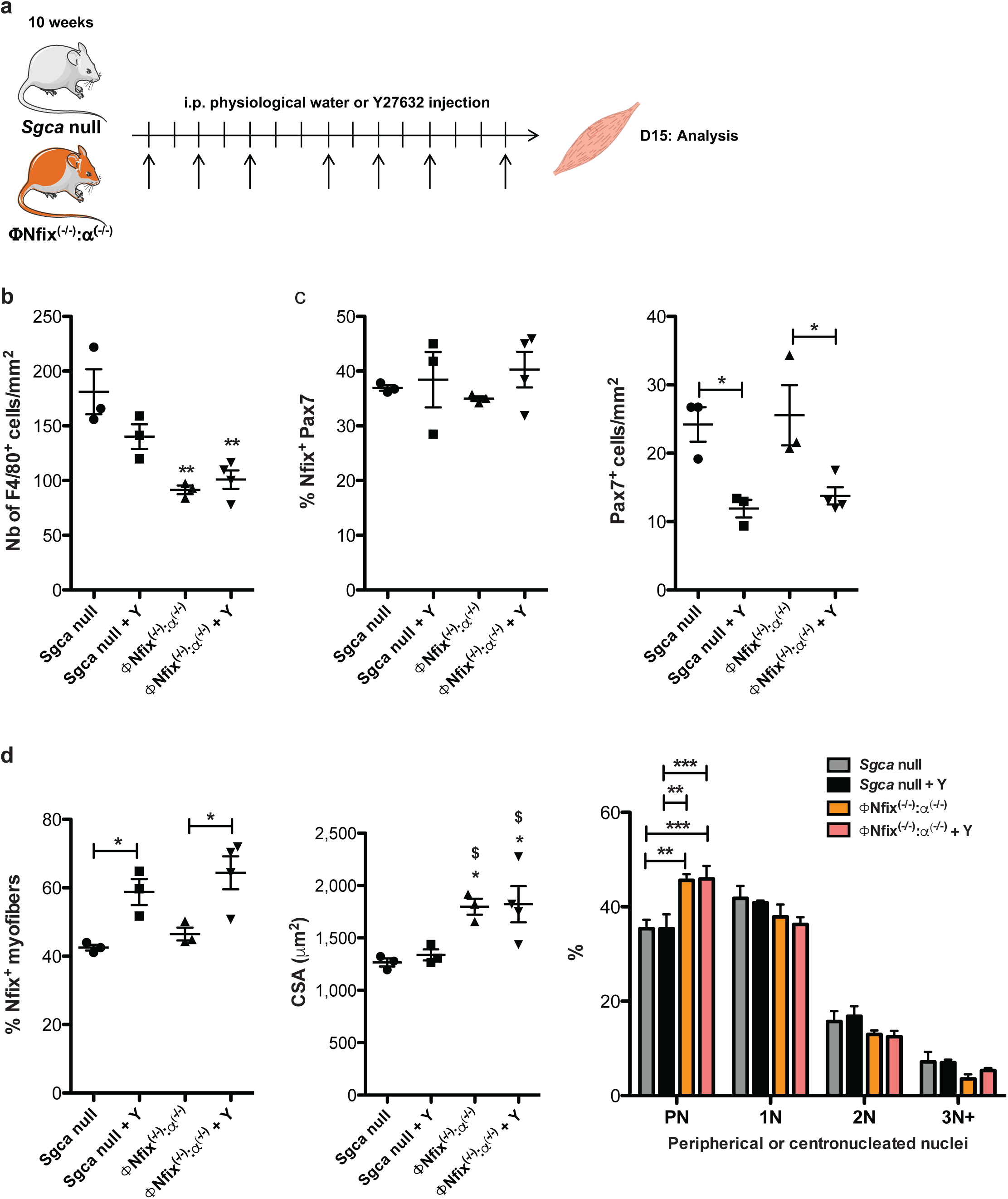
Pharmacological ROCK1 inhibition strategy, number of MPs and Nfix^+^ SCs. a) 10 weeks old *Sgca* null and ΦNfix^(-/-)^:α^(-/-)^ mice were i.p. injected 3 times per week for 2 weeks with physiological water or Y27632 ROCK inhibitor and analyzed. b) Number of F4/80^+^ cells per mm^2^ of *Sgca* null and ΦNfix^(-/-)^:α^(-/-)^ TA muscles of mice i.p. injected with physiological water or Y. c) Percentage of double Nfix^+^/Pax7^+^ cells and number of Pax7^+^ cells/mm^2^ of *Sgca* null and ΦNfix^(-/-)^:α^(-/-)^ TA muscles of mice i.p. injected with physiological water or Y. d) Percentage of Nfix^+^ myofibers, CSA and percentage of perinucleated and centronucleated (from 1 to 3^+^ nuclei) myofibers of *Sgca* null and ΦNfix^(-/-)^:α^(-/-)^ TA muscles of mice i.p. injected with physiological water or Y. Statistical significance was determined using one-way Anova test or two-way Anova test.* vs *Sgca* null: * p<0,05 ; ** p<0,01. $ vs *Sgca* null + Y: $ p<0,05. Results are means ± SEM of at least three independent experiments. Scale bar = 50μm.

**Fig S4.**
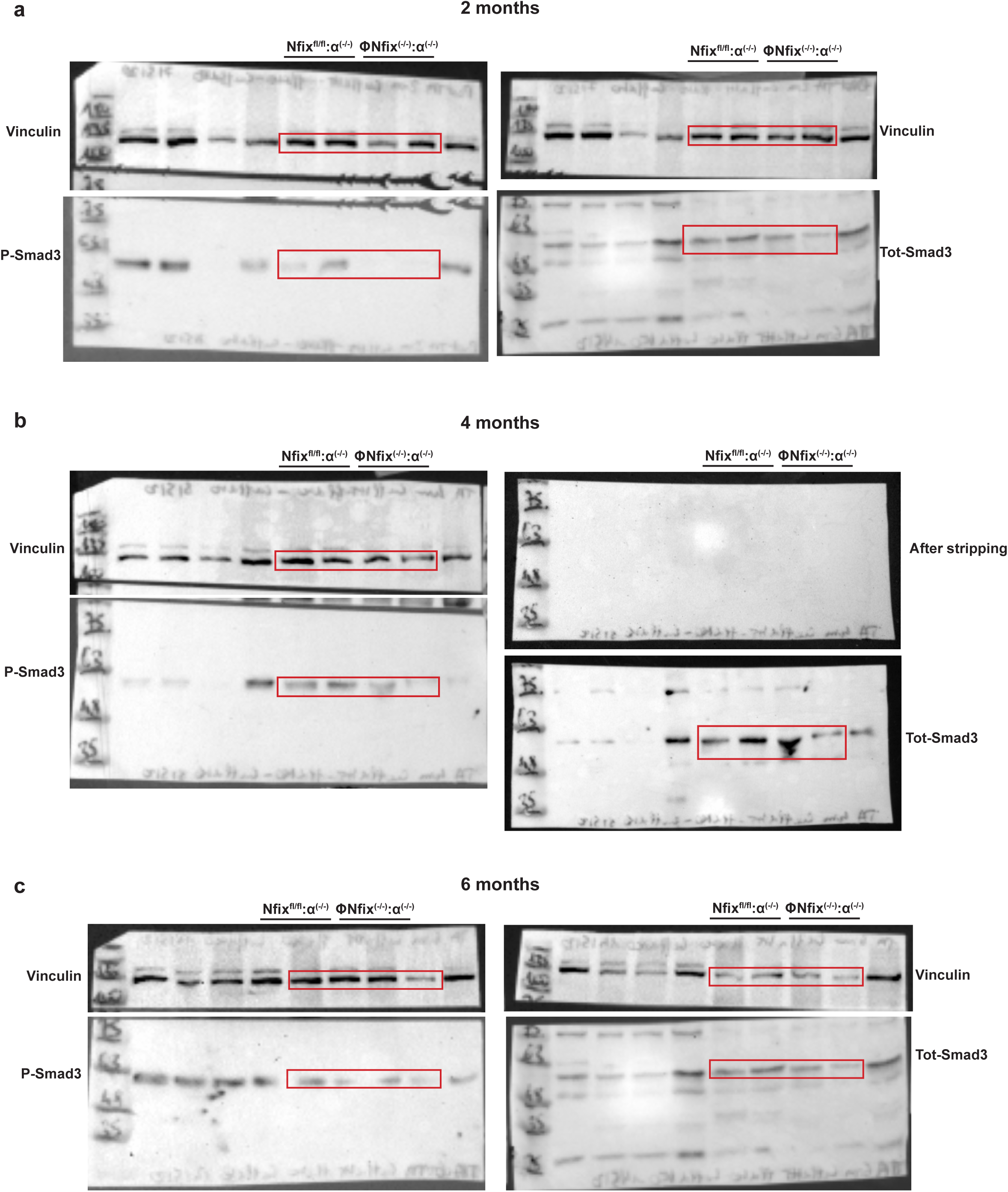

